# Environmental risk reshapes high-confidence schizophrenia risk genes during a developmental window of vulnerability

**DOI:** 10.64898/2026.10.27.762379

**Authors:** Alison Brack, Rhushikesh A. Phadke, Luke Fournier, Matthew J. Boberg, Maria Salgado, Alberto Cruz-Martín

## Abstract

Rare coding variants and environmental exposures such as early-life stress contribute to schizophrenia risk, but whether environmental adversity alters the expression of genetically implicated genes remains unclear. Combining the limited bedding and nesting (LBN) model of early-life stress (ELS) with 24-plex single-cell mFISH registered to the Allen Mouse Brain Atlas, we profiled the ten genes identified in the original SCHEMA study, the complement gene *C4b* and activity markers in 1.29 million cells across four postnatal ages. Whole-brain risk-gene upregulation was greatest at postnatal day 21 (P21), with region- and cell-type-selective effects that prominently involved the hypothalamus. *Sp4* and *Cacna1g* showed distinct temporal responses: *Sp4* increased at P21 and decreased at P90 in hypothalamic oligodendrocytes, whereas *Cacna1g* decreased at P10. ELS-associated *C4b* upregulation extended across neuronal and glial populations, including oligodendrocytes and OPCs. Across regional cell types, P10 *C4b* effects were positively correlated with P21 *Grin2a* effects. These findings suggest that ELS perturbs the developmental expression of genes implicated in schizophrenia, identifying potential molecular links between environmental adversity and genetic vulnerability.

## Introduction

Schizophrenia (SCZ) is a debilitating neurodevelopmental disorder characterized by positive symptoms, such as hallucinations and delusions, as well as disabling negative and cognitive symptoms^1^. The disorder typically emerges in late adolescence and early adulthood^1^. Social isolation and difficulty managing daily life are common challenges. The lifetime risk of suicide is approximately 5%, with heightened risk early in the course of illness^2^. Available antipsychotic medications primarily target positive symptoms but have limited capacity to ameliorate negative and cognitive symptoms^3,4^.

In 2022, the Schizophrenia Exome Sequencing Meta-Analysis (SCHEMA) study analyzed exome data from over 100,000 participants and identified ten genes in which ultra-rare coding variants were associated with SCZ at exome-wide significance^5^. Associated variant classes included protein-truncating and predicted damaging missense variants, with estimated odds ratios ranging from approximately 3 to 50. These findings substantially expanded the set of genes robustly implicated through rare coding variation, opening new avenues for research into SCZ pathogenesis. The diversity of these genes, which span synaptic function, transcriptional regulation and protein ubiquitination, suggests that multiple biological processes contribute to SCZ susceptibility. However, the spatiotemporal expression patterns of these genes during postnatal development, and how their expression responds to early-life stress (ELS), remain incompletely characterized.

Monozygotic twins of individuals with SCZ show incomplete concordance for the disorder, indicating that genetic susceptibility is not fully deterministic and that environmental and other non-genetic factors contribute to its development^6,7^. During childhood and adolescence, exposure to ELS, including abuse, neglect and bullying, is associated with approximately 2.8-fold higher odds of psychosis^8,9^. Childhood maltreatment has also been associated with elevated inflammatory markers in adulthood, suggesting that changes in immune regulation may contribute to the long-term consequences of ELS^10^. Experimental studies in mice further suggest that prenatal immune activation and subsequent peripubertal stress can interact to alter adult behavior and neurochemistry^11^. In rodents, ELS alters hypothalamic-pituitary-adrenal (HPA) axis function and oligodendrocyte maturation^12,13^.

How early-life stress reshapes the expression of the ten genes implicated by SCHEMA across brain regions, cell types and postnatal development remains incompletely understood. To address this gap, we combined the limited bedding and nesting (LBN) model of ELS with 24-plex single-cell multiplexed fluorescence *in situ* hybridization (mFISH) across four postnatal ages. We profiled the mouse orthologs of the ten SCHEMA genes, the complement gene *C4b*, three additional neuronal markers and ten cell-type markers, registering tissue images to the Allen Mouse Brain Atlas. We included *C4b* based on evidence linking genetically predicted increases in human brain *C4A* expression to SCZ risk^14^ and showing that increased mouse *C4* expression disrupts prefrontal cortical connectivity and reduces social interaction^15^. We find that ELS produces a response that is most pronounced at P21, is regionally and cell-type selective, and prominently involves the hypothalamus. *Sp4* and *Cacna1g* define two distinct modes of this response: a P21 increase in *Sp4* that reverses by adulthood within the same defined oligodendrocyte subtype, and an early *Cacna1g* decrease that is already present at P10. Finally, region–cell-type groups with larger *C4b* changes at P10 tend to show larger *Grin2a* changes at P21. These data provide a developmental, cell-type-resolved map of how ELS alters the expression of genes implicated in SCZ risk.

## Results

### A single-cell mFISH atlas of SCHEMA risk-gene expression across postnatal development

To characterize SCHEMA risk-gene expression across postnatal development and following ELS, we collected whole brains from mice exposed to LBN from P2 to P9 and controls at four ages: shortly after exposure (P10), at weaning (P21), during adolescence (P40), and in adulthood (P90) (Fig. 1a).

**Fig. 1.**
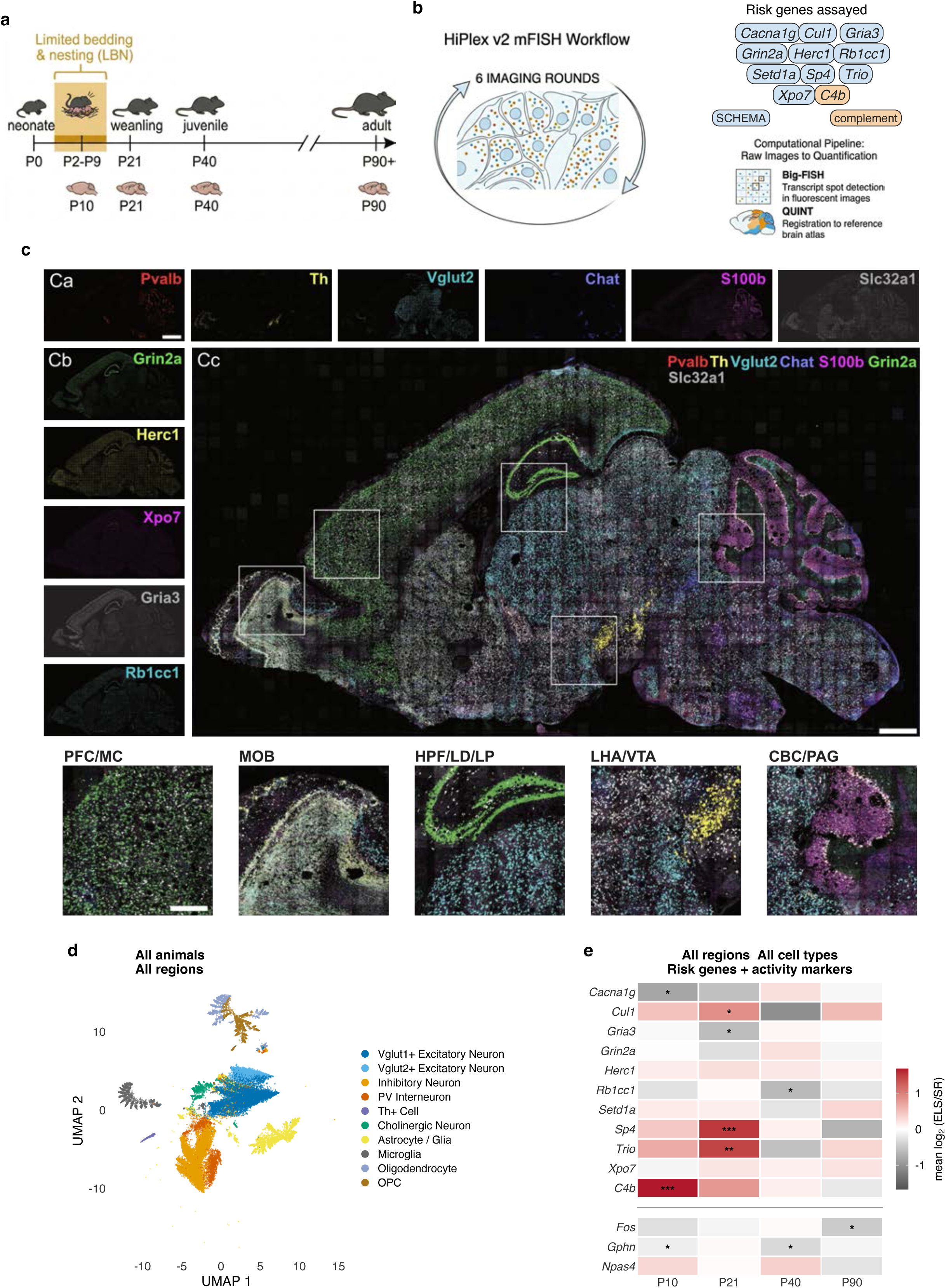
Whole-brain single-cell mFISH atlas of SCHEMA risk-gene expression across postnatal development. a, Limited bedding and nesting (LBN) paradigm used to model early-life stress (ELS) and sampling timeline. SR, standard-reared. b, HiPlex v2 mFISH workflow, risk panel (ten SCHEMA genes, blue; complement gene *C4b*, amber) and analysis pipeline. c, Representative sagittal section with single-channel images of cell-type markers (Ca) and SCHEMA genes (Cb), merged composite (Cc) and regional crops of the boxed areas: PFC/MC, prefrontal and motor cortex; MOB, main olfactory bulb; HPF/LD/LP, hippocampal formation and lateral dorsal and lateral posterior thalamic nuclei; LHA/VTA, lateral hypothalamic area and ventral tegmental area; CBC/PAG, cerebellar commissure and periaqueductal gray. Scale bars, 2 mm (Ca, Cb), 750 µm (Cc) and 300 µm (crops). d, UMAP of classified cells colored by cell type (30,000 of the 1,163,386 classified cells with at least 6 of 24 genes detected); cell types were assigned from marker expression before embedding (Methods). e, Mean litter-adjusted log₂(ELS/SR) for each gene at each age across all region × cell type strata; activity and inhibitory-scaffold markers below the rule. Asterisks, whole-brain contrast at that age (litter-adjusted mixed model, Benjamini–Hochberg adjusted across the 44 risk-gene × age tests, or within each activity marker across ages). n = 4 animals per condition per age (32 animals, 10 litters). *P < 0.05, **P < 0.01, ***P < 0.001.

To profile these genes across ages and rearing conditions, we built a 24-gene HiPlex panel comprising the ten SCHEMA genes^5^, the well-established SCZ risk gene complement component 4 (*C4b*)^14,15^, three activity and inhibitory-scaffold markers (*Fos*, *Npas4* and *Gphn*) and ten cell-type markers (Fig. 1b). We imaged whole sagittal sections from 32 mice (4 SR and 4 ELS at each of P10, P21, P40 and P90, both sexes, from 10 litters), detected puncta with Big-FISH^16^ and registered each cell to the Allen Mouse Brain Atlas with QUINT^17^ (Fig. 1b,c). Of 1,907,152 segmented cells, 1,288,350 (67.6%) met marker-based criteria and were assigned to one of ten cell types across ten brain regions (Fig. 1d). In a UMAP embedding of 21 panel genes (the cell-type markers, SCHEMA genes and *C4b*), these populations occupied distinct territories (Fig. 1d). Across the whole brain, ELS altered expression of the risk panel mainly at P10 and P21 (Fig. 1e). At P10, *C4b* was increased (log₂ fold change (log₂FC) = 1.67) and *Cacna1g* decreased (log₂FC = −0.78); at P21, *Sp4* (1.48), *Trio* (1.97) and *Cul1* (1.41) were increased and *Gria3* (−0.57) decreased; at P40, only *Rb1cc1* was changed (−0.71); and at P90, no risk gene was significantly changed (whole-brain litter-adjusted mixed models, Benjamini–Hochberg (BH) correction across 11 genes × 4 ages; asterisks in Fig. 1e). Under standard rearing, normalized whole-brain *C4b* expression increased from P10 to P40, whereas *Cacna1g* expression declined after P21. *Trio* expression was lowest at P21, coinciding with the age at which ELS increased its expression (Supplementary Fig. 1a). Among the activity and scaffold markers, *Gphn* was modestly reduced at P10 and P40 and *Fos* at P90, whereas *Npas4* was unchanged at every age (Fig. 1e). Together, these findings established a single-cell atlas of postnatal risk-gene expression and identified P10 and P21 as the principal ages at which ELS altered whole-brain expression.

### ELS-associated risk-gene upregulation was greatest at P21 and varied across regions and cell types

Whole-brain risk-gene upregulation was greatest at P21 (Fig. 2a,b). The signed mean log₂FC across the eleven risk-panel genes was 0.15 at P10, 0.42 at P21, −0.06 at P40 and 0.02 at P90 (Fig. 2b), and the condition × age interaction was significant for *Sp4*, *C4b*, *Cul1*, *Cacna1g* and *Trio* (likelihood-ratio tests, BH-adjusted P ≤ 0.032), indicating that the effect of ELS on these genes depended on age. Interestingly, *Sp4*, which was increased at P21, showed a nonsignificant decrease at P90 (log₂FC = −0.65, adjusted P = 0.073). Under standard rearing, normalized whole-brain *Sp4* expression increased from its lowest observed level at P21 to its highest at P90 (Supplementary Fig. 1a). When we resolved the response by region and by cell type with ages pooled, we found that it was selective (Fig. 2c). The most consistent stratum-level change was a reduction of *Rb1cc1*, significant in six of nine regions and nine of ten cell types. *C4b*, *Trio* and *Sp4* were increased in most regions and cell types, but with ages pooled, few of these increases survived correction, consistent with the age-dependent effects described above. Comparing gene-level effects across ages separated two components of the response (Fig. 2d). *C4b* and *Cacna1g* were already changed at P10, whereas *Sp4*, *Trio*, *Cul1* and *Gria3* were changed only at P21. Later effects were smaller (Fig. 2e): *Rb1cc1* was the only gene changed at P40, and no gene was significantly changed at P90. At the cellular level, *C4b* was increased in oligodendrocytes of the fiber tracts at P10 (P = 0.041) and *Trio* in OPCs of the hypothalamus at P21 (P = 0.027; per-animal linear mixed models with litter as a random effect; Fig. 2f,g). Together, these results showed that ELS reshapes SCZ-risk gene expression in a developmentally timed, bidirectional manner that peaks at P21.

**Fig. 2.**
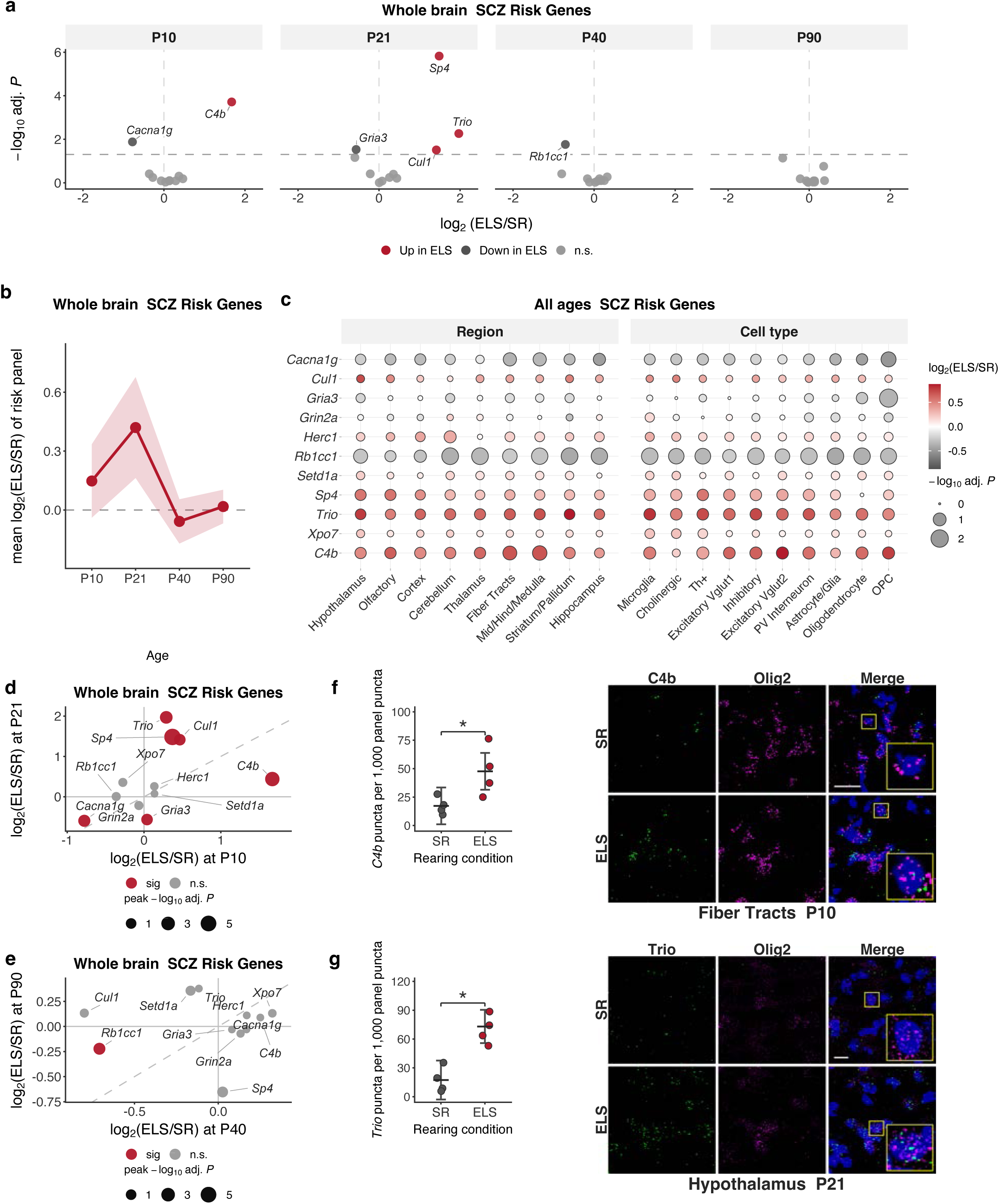
ELS produces a P21-peaking, regionally and cell-type-selective response concentrated on SCHEMA risk genes. a, Volcano plots of ELS versus SR at each age for the 11 risk-panel genes; significant genes are labeled. Dashed line, adjusted P = 0.05 (Benjamini–Hochberg across 11 genes × 4 ages). b, Mean log₂(ELS/SR) across the 11 risk-panel genes at each age; shading, s.e.m. across genes. c, Effects by region (left) and cell type (right), pooling ages. Fill, log₂(ELS/SR); size, −log₁₀ adjusted P. d,e, Gene-level log₂(ELS/SR) at P10 versus P21 (d) and P40 versus P90 (e); colored points, significant at either age; dashed line, y = x. f, *C4b* in oligodendrocytes of the fiber tracts at P10. Left, per-animal expression (puncta per 1,000 panel puncta); bars, linear mixed model estimate with litter as a random effect; whiskers, 95% confidence interval; bracket, ELS versus SR (n = 4 SR, 4 ELS; 4 litters; P = 0.041). Right, target gene (green), cell-type marker (magenta) and DAPI (blue); SR above, ELS below; callouts, 3× magnification of the cell in the yellow box. Scale bar, 18.9 µm. g, *Trio* in *O*PCs of the hypothalamus at P21, as in f (n = 4 SR, 4 ELS; 5 litters; P = 0.027). Scale bar, 10.1 µm. n = 4 animals per condition per age (32 animals, 10 litters) unless noted.

### The hypothalamus is a convergent site of ELS vulnerability

At P21, *Sp4*, *C4b* and *Cul1* were increased in most regions, with the largest *Sp4* (log₂FC = 2.26) and *Cul1* (1.67) effects observed in the hypothalamus (Fig. 3a). These findings, together with the hypothalamus’s central role in regulating the stress response through the hypothalamic-pituitary-adrenal (HPA) axis^18^, prompted us to examine this region in greater detail.

**Fig. 3.**
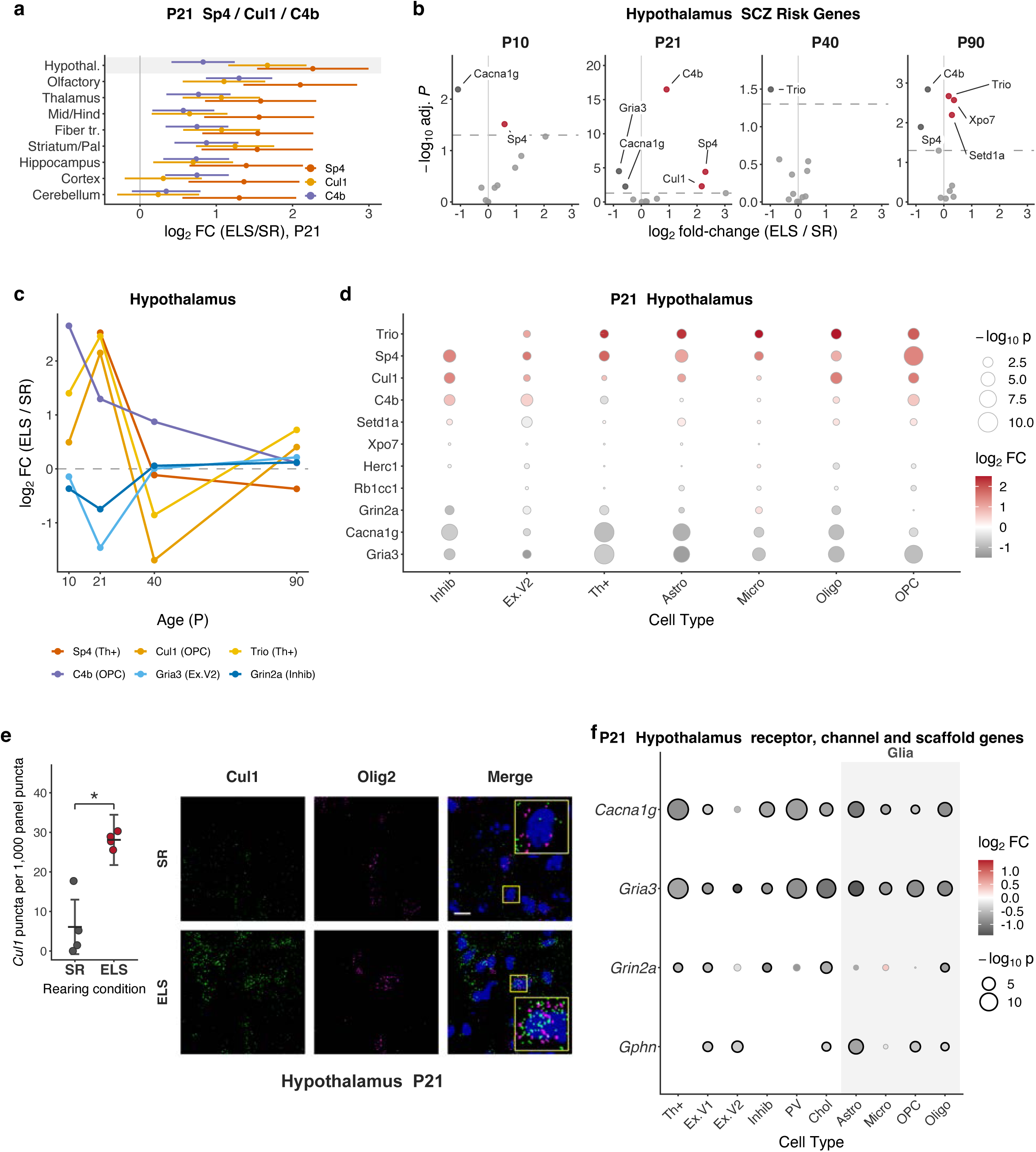
The hypothalamus is a convergent site of ELS vulnerability. a, log₂(ELS/SR) at P21 for *Sp4*, *Cul1* and *C4b* by region (mean ± 95% confidence interval); hypothalamus shaded. b, Volcano plots of risk-panel genes in the hypothalamus at each age, as in Fig. 2a (Benjamini–Hochberg across 11 genes × 4 ages); y axes differ between ages. c, log₂(ELS/SR) trajectories of six hypothalamic gene × cell type pairs. d, P21 hypothalamic response by cell type and gene after correction for optical spillover from neighboring cells. Fill, log₂(ELS/SR); size, −log₁₀ P; blank, model did not converge. e, *Cul1* in OPCs of the hypothalamus at P21, as in Fig. 2f (n = 4 SR, 4 ELS; 5 litters; P = 0.017). Scale bar, 10.1 µm. f, Receptor, channel and scaffold genes in neuronal and glial (shaded) populations of the P21 hypothalamus, as in d; black outline, adjusted P < 0.05. n = 4 animals per condition per age (32 animals, 10 litters) unless noted.

Within the hypothalamus, ELS changed *Cacna1g* and *Sp4* at P10, and the response peaked at P21 with *C4b* (log₂FC = 0.90), *Sp4* (2.30) and *Cul1* (2.17) increased and *Gria3* (−0.81) and *Cacna1g* (−0.58) decreased (Fig. 3b). Unlike the whole brain, the hypothalamus continued to show significant ELS-associated differences in adulthood: at P90, *C4b* (−0.58) and *Sp4* (−0.83) were decreased and *Trio*, *Xpo7* and *Setd1a* were increased (Fig. 3b). Selected gene–cell-type combinations showed distinct trajectories of ELS-associated expression differences across ages (Fig. 3c). To determine which hypothalamic cell types carried the P21 response, and to control for optical spillover of puncta from neighboring cells, we refit the models after correcting each cell for its local neighborhood.

Using this approach, we showed that the P21 response was distributed across neuronal and glial populations (Fig. 3d): *Sp4* was increased in all seven displayed cell types, *Cul1* most strongly in OPCs, oligodendrocytes and inhibitory neurons, *C4b* in inhibitory neurons, OPCs and excitatory neurons, and *Trio* in glial populations and Th⁺ cells. Under standard rearing, normalized *Trio* expression in hypothalamic Th⁺ cells was lowest at P21 and highest at P40 (Supplementary Fig. 2c). *Cul1* was increased in hypothalamic OPCs at the animal level (P = 0.017; Fig. 3e).

Receptor and channel genes were reduced across the same populations (Fig. 3f): *Gria3* and *Cacna1g* were decreased in astrocytes, oligodendrocytes and OPCs as well as in neurons; *Grin2a* in inhibitory, cholinergic, Th⁺ and *Vglut1*⁺ neurons and oligodendrocytes, and the inhibitory postsynaptic scaffold *Gphn* in astrocytes, OPCs, oligodendrocytes and excitatory and cholinergic neurons. Together, these findings indicated a coordinated hypothalamic ELS response that peaks at P21, spans neuronal and glial cell types and persists, in altered form, into adulthood.

### *Sp4* and *Cacna1g* define two distinct modes of SCHEMA vulnerability

Ordering the risk-panel genes by the difference between their P21 and P90 effects placed *Sp4* at the top, and *Cacna1g* showed the largest P10 decrease (Fig. 4a). At the whole-brain level, *Sp4* was increased at P21 and trended below standard-reared levels at P90, whereas *Cacna1g* was decreased at P10, showed a nonsignificant decrease at P21 and showed no significant difference at P40 (Fig. 4b). The P10 *Cacna1g* decrease was broad across regions and cell types and was no longer significant at P40 (Fig. 4c). The P21 *Sp4* increase was likewise broad, and by P90 many of the same strata showed decreases (Fig. 4d). Within PV⁺ interneurons, *Cacna1g* was lower in every region at P10, significantly so in five of nine regions and most strongly in the hypothalamus, with no significant reductions detected at P40 (Fig. 4e). Within oligodendrocytes, *Sp4* was increased at P21 and decreased at P90 in all six regions sampled from at least two litters per condition (Fig. 4f). Per-animal quantification in hypothalamic oligodendrocytes confirmed the *Sp4* reversal within a single population, with an increase at P21 (P = 0.037) and a decrease at P90 (P = 0.025; Fig. 4h). Under standard rearing, normalized *Sp4* expression in hypothalamic oligodendrocytes was lowest at P21 and increased through P90 (Supplementary Fig. 2c). In hypothalamic PV⁺ interneurons, *Cacna1g* expression was lower following ELS at P10, although the animal-level comparison did not reach significance (P = 0.052); no significant difference was detected at P40 (P = 0.20; Fig. 4g). Under standard rearing, normalized *Cacna1g* expression in hypothalamic PV⁺ neurons declined after P21 (Supplementary Fig. 2c). These two genes therefore highlighted contrasting patterns of the SCHEMA response to ELS: a broad *Sp4* increase at P21 followed by decreases in adulthood, and an early *Cacna1g* decrease that was no longer significant at P40.

**Fig. 4.**
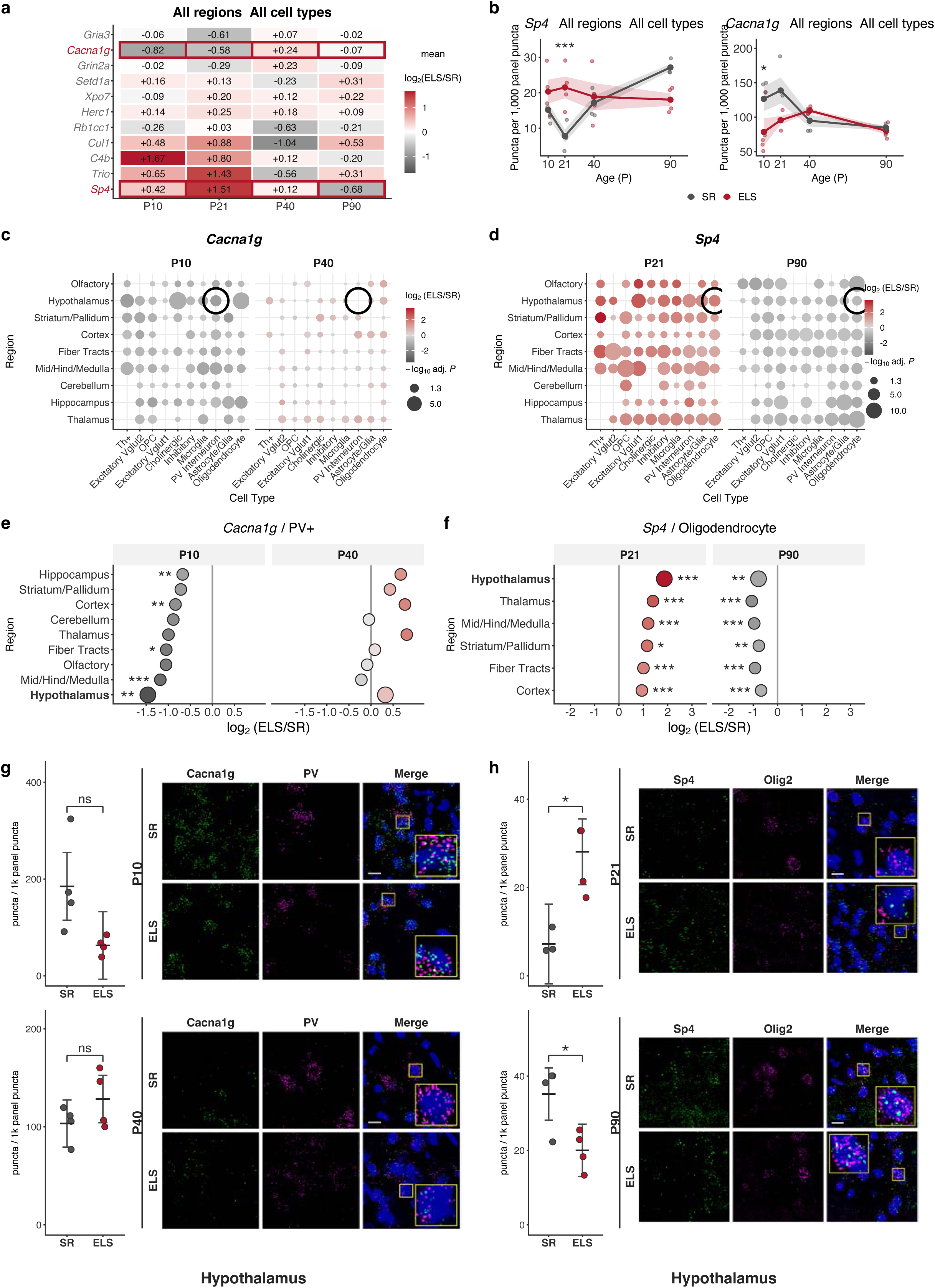
*Sp4* and *Cacna1g* define two distinct modes of SCHEMA vulnerability. a, Mean log₂(ELS/SR) for each risk-panel gene at each age, ordered by the P21 minus P90 effect; *Sp4* and *Cacna1g* outlined. b, Whole-brain *Sp4* and *Cacna1g* expression (puncta per 1,000 panel puncta, all classified cells) by age. Lines and shading, mean ± s.e.m.; dots, individual animals; asterisks, whole- brain ELS versus SR contrast at that age, as in Fig. 1e. c,d, *Cacna1g* at P10 and P40 (c) and *Sp4* at P21 and P90 (d) by region and cell type. Fill, log₂(ELS/SR); size, −log₁₀ adjusted P; ring, stratum quantified in g (c) or h (d); blank, stratum sampled in fewer than two animals per condition or without a model P value. e, *Cacna1g* in PV⁺ interneurons by region at P10 and P40, for regions sampled from at least two litters per condition at both ages; hypothalamus in bold; asterisks, Benjamini–Hochberg- adjusted P across strata for each gene and age. f, *Sp4* in oligodendrocytes by region at P21 and P90, as in e. g, *Cacna1g* in hypothalamic PV⁺ interneurons at P10 (top; n = 4 SR, 4 ELS; 4 litters; P = 0.052) and P40 (bottom; n = 4 SR, 4 ELS; 5 litters; P = 0.20), as in Fig. 2f. h, *Sp4* in hypothalamic oligodendrocytes at P21 (top; n = 3 SR, 4 ELS; 5 litters; P = 0.037) and P90 (bottom; n = 4 SR, 4 ELS; 6 litters; P = 0.025), as in Fig. 2f. Scale bars, 10.1 µm. n = 4 animals per condition per age (32 animals, 10 litters) unless noted.

### Early ELS effects were associated with P21 expression changes across regional cell types

Because ELS effects were prominent at P10 and P21, we next asked whether expression changes at these ages were correlated across regional cell types. For each of the 14 panel genes, we estimated ELS effects within each region × cell-type stratum at P10 and P21, then calculated cross-age Spearman correlations for every gene pair across matched strata (Fig. 5a). Across regional cell types, P10 *C4b* effects were positively correlated with P21 *Grin2a* (Spearman ρ = 0.49, adjusted P = 2.7 × 10⁻⁴) and *Cacna1g* effects (ρ = 0.44, adjusted P = 0.0023; Fig. 5b,e). P10 *Cacna1g* effects were positively correlated with P21 *Fos* effects (ρ = 0.31, adjusted P = 0.047; Fig. 5c,f). Illustrating these cross-age associations, *C4b* was increased at P10 (P = 0.011) and *Grin2a* at P21 (P = 0.037) in OPCs of the mid/hindbrain and medulla (Fig. 5h). In hypothalamic cholinergic neurons, *Cacna1g* was decreased at P10 (P = 0.0040) and *Fos* at P21 (P = 0.032; Fig. 5i).

**Fig. 5.**
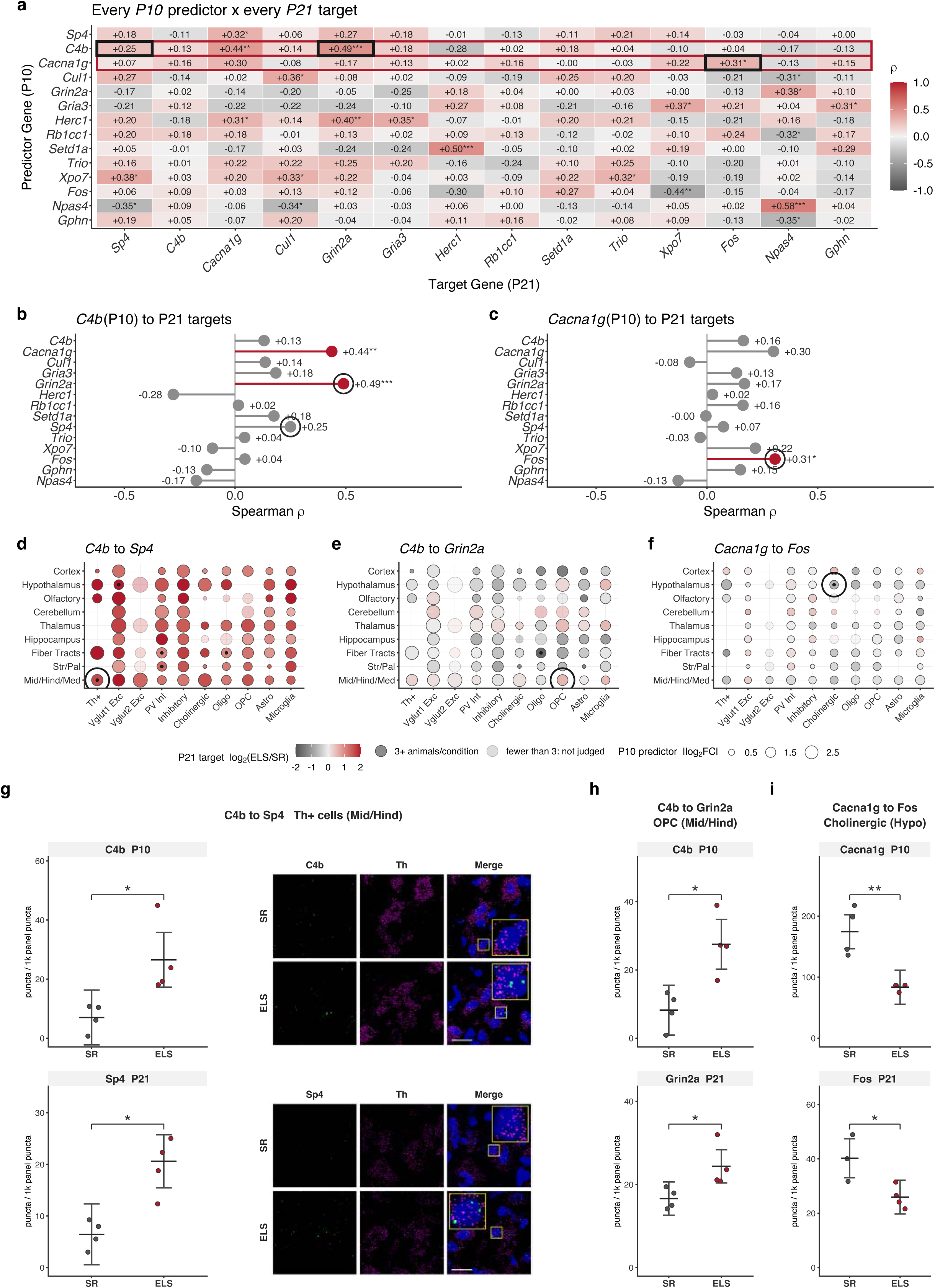
P10 ELS effects are associated with distinct P21 outcomes in the same regional cell types. a, Spearman correlations across region × cell type strata between the P10 effect of each predictor gene and the P21 effect of each target gene (log₂(ELS/SR); Benjamini–Hochberg across 196 pairs). Red outlines, predictor rows shown in b,c; black outlines, pairs shown in d–f; asterisks, adjusted *P < 0.05, **P < 0.01, ***P < 0.001. Associations do not imply causation. b,c, *C4b* (b) and *Cacna1g* (c) rows of a; red, adjusted P < 0.05; rings, targets shown in d–f. d–f, Strata for *C4b* to *Sp4* (d; ρ = +0.25, 79 strata, adjusted P = 0.15), *C4b* to *Grin2a* (e; ρ = +0.49, 80 strata, adjusted P = 2.7 × 10⁻⁴) and *Cacna1g* to *Fos* (f; ρ = +0.31, 81 strata, adjusted P = 0.047). Fill, P21 target stratum effect (log₂(ELS/SR)); size, magnitude of the P10 predictor stratum effect (the values correlated in a); faded, fewer than three animals per condition; center dot, both effects adjusted P < 0.05; ring, stratum quantified in g–i. g–i, Per-animal quantification of the P10 predictor (top) and P21 target (bottom) in one stratum, as in Fig. 2f. g, *C4b* to *Sp4* in Th⁺ cells of the mid/hindbrain and medulla (*C4b*: n = 4 SR, 4 ELS; 4 litters; P = 0.027. *Sp4*: n = 4 SR, 4 ELS; 5 litters; P = 0.041). Scale bar, 18.9 µm. h, *C4b* to *Grin2a* in OPCs of the mid/hindbrain and medulla (*C4b*: n = 4 SR, 4 ELS; 4 litters; P = 0.011. *Grin2a*: n = 4 SR, 4 ELS; 5 litters; P = 0.037). i, *Cacna1g* to *Fos* in cholinergic neurons of the hypothalamus (*Cacna1g*: n = 4 SR, 4 ELS; 4 litters; P = 0.0040. *Fos*: n = 3 SR, 4 ELS; 5 litters; P = 0.032). n = 4 animals per condition per age (32 animals, 10 litters) unless noted.

The association between P10 *C4b* and P21 *Sp4* effects across strata was positive but did not survive correction (ρ = 0.25, adjusted P = 0.15; Fig. 5d). In Th⁺ cells of the mid/hindbrain and medulla, however, both *C4b* at P10 (P = 0.027) and *Sp4* at P21 (P = 0.041) were increased (Fig. 5g). Additional cross-age associations survived correction outside the *C4b* and *Cacna1g* comparisons highlighted here (Fig. 5a). These associations linked ELS effects across matched regional cell types in different animals; they did not establish that early changes caused later ones, but identified candidate relationships for future mechanistic studies.

## Discussion

Using a whole-brain, single-cell mFISH dataset spanning four postnatal ages, we found that ELS altered SCHEMA risk-gene and *C4b* expression shortly after exposure, with region-specific differences still detectable in adulthood. Whole-brain risk-gene upregulation was greatest at P21, while region- and cell-type-resolved analyses identified the hypothalamus as a prominent site of the response, involving both neuronal and glial populations, including the oligodendrocyte lineage. Within this response, *Sp4* showed widespread upregulation at P21 followed by reductions in adulthood, whereas *Cacna1g* showed an early decrease that was no longer significant at P40. Across regional cell types, P10 *C4b* effects were positively correlated with P21 *Grin2a* effects.

The concentration of significant changes at P10 and P21 places the ELS response in a juvenile window that precedes adolescence, a developmental period relevant for SCZ^19^ given that adolescence and early adulthood are the typical ages of onset^1^. These early expression changes may contribute to later vulnerability, even though we detected no significant whole-brain differences in risk-gene expression at P90. Whether they contribute to the long-lasting consequences of ELS documented in mice^12^ and humans^20^ remains to be tested.

Although not among the SCHEMA genes, mouse *C4b* was included in our panel because of the association between genetic variation at the human complement component 4 (*C4*) locus and SCZ risk^14^. *C4b* showed the strongest whole-brain upregulation at P10, the first age examined; no significant whole-brain difference was detected at P40. Although astrocytes have been investigated as a source of secreted C4 and a potential therapeutic target^21^, ELS-associated *C4b* upregulation extended across neuronal and glial populations, including fiber-tract oligodendrocytes at P10 and hypothalamic inhibitory neurons, excitatory neurons and OPCs at P21. These findings argue for considering multiple cellular sources when investigating stress-associated complement dysregulation.

Human *C4* structural alleles associated with higher brain *C4A* expression confer greater SCZ risk^14^. Experimentally, developmental *C4* overexpression in mouse prefrontal cortex reduced dendritic spine density and excitatory synaptic drive and impaired social interaction^15^, providing a potential link to the reduced cortical spine density reported in postmortem SCZ tissue^22^. Notably, *C4* expression was no longer elevated at P60, yet social deficits persisted into adulthood^15^, suggesting that transient developmental elevations can have lasting behavioral consequences. Across regional cell types in our study, P10 *C4b* effects were positively correlated with P21 *Grin2a* and *Cacna1g* effects. These associations linked early and later expression changes across matched populations in different animals; they did not establish that one drove the other. The association between P10 *C4b* and P21 *Sp4* effects was positive but did not survive correction, leaving this relationship unresolved.

*Sp4* and *Cacna1g* showed distinct temporal responses to ELS. *Sp4* was increased at P21 across regions and cell types, with a reversal to reduced expression at P90 observed in hypothalamic oligodendrocytes. *Cacna1g*, by contrast, was reduced at P10, with no significant whole-brain difference detected at P40. The early increase in *Sp4* expression contrasted with the loss-of-function variants associated with SCZ risk, whereas the decrease in *Cacna1g* expression was directionally consistent with reduced gene function. Whether the early *Cacna1g* decrease or the later *Sp4* decrease produced functional consequences resembling those of SCZ-associated loss-of-function variants remains to be tested. Because each animal was sampled at a single age, these trajectories describe population-level rather than within-animal change.

A recent SCHEMA 2.0 preprint retained exome-wide significant associations for *SP4*, *CUL1*, *SETD1A*, *HERC1*, *RB1CC1* and *XPO7*, whereas *CACNA1G*, *GRIN2A* and *GRIA3* did not meet the updated 5% FDR threshold^23^. Our panel was based on the original SCHEMA findings, but evidence from other approaches supports continued investigation of these genes. *GRIN2A* was independently prioritized through common-variant schizophrenia GWAS fine-mapping ^24^, while reduced *GRIA3* expression was reported in postmortem schizophrenia samples and patient-derived cerebral organoids^25^. Experimental evidence also linked *Cacna1g* loss to SCZ-relevant sleep and network abnormalities^26^, although its genetic association with SCZ remains uncertain. These findings emphasize the importance of considering genetic association, disease-associated expression and experimental function as complementary but distinct forms of evidence.

Our identification of *Sp4* as an upregulated gene after ELS is perhaps counterintuitive, since loss-of-function mutations in *SP4* increase SCZ risk^5^. An optimal range of *Sp4* expression may be required to maintain appropriate expression of the synaptic genes it regulates^27^. SCZ risk genes, including other SCHEMA genes, are enriched among potential *SP4* targets^28^. Interestingly, the biphasic *Sp4* response may reflect a loss of developmental regulation rather than a transient pulse: under standard rearing, *Sp4* expression changes dynamically across development, whereas under ELS it remains near a constant level from P10 through adulthood. One possibility is that ELS blunts the normal age-dependent regulation of *Sp4* expression.

The hypothalamus was a convergent site of the ELS response. ELS has been shown to lead to HPA axis dysregulation^29^, and chronic HPA axis activation leads to the overproduction of glucocorticoids, which can have detrimental effects on brain health. HPA activity is increased in individuals with SCZ^18^. The hypothalamic changes we observe may therefore reflect altered HPA axis activity, although we did not measure HPA function directly. The hypothalamic response was not limited to neurons: at P21 it included astrocytes, oligodendrocytes and OPCs, and receptor and channel genes such as *Gria3* and *Cacna1g* were reduced in glial as well as neuronal populations.

Changes in myelin have long been observed in SCZ^30,31^, and juvenile social isolation, a related mouse model of ELS, impairs oligodendrocyte maturation and myelination in the medial prefrontal cortex between P21 and P35^32^. Interestingly, the induction of SCHEMA genes in hypothalamic oligodendrocyte-lineage cells at P21 falls within this same window. In the hypothalamus, ELS-associated expression differences remained detectable at P90, with *C4b* and *Sp4* reduced and *Trio, Xpo7* and *Setd1a* increased.

The persistence of regional differences despite the absence of significant whole-brain effects at P90 highlighted the importance of spatially and cellularly resolved measurements. Whole-brain averages can obscure changes restricted to particular populations or combine effects of opposing direction. Thus, the lack of a detectable whole-brain difference in adulthood should not be interpreted as complete normalization of the ELS response.

A meta-analysis of genetic mouse models carrying loss-of-function mutations in SCHEMA- associated genes identified age- and region-dependent transcriptomic changes, with convergence in the striatum and thalamus^33^. Although the hypothalamus was not examined, these findings complement our results by emphasizing that the consequences of perturbing SCZ-associated genes depend on anatomical and developmental context.

*Cacna1g* was reduced in PV⁺ interneurons at P10 across regions. PV⁺ interneurons are central to SCZ models that implicate NMDA receptor hypofunction^34^ and the resulting hypoactivity of PV neurons^35^, and to the oscillatory activity that coordinates information flow^36^. A recent preprint further linked *Cacna1g* loss to disrupted sleep spindles, slow oscillations and cortical synchrony^26^, supporting a role for this channel in network coordination. The decrease in *Cacna1g* expression within hypothalamic PV⁺ interneurons did not reach significance in the per-animal analysis (P = 0.052), warranting confirmation in a larger cohort. Strata with larger P10 *Cacna1g* changes also tended to show larger P21 *Fos* changes.

This study had several limitations. First, although ELS was experimentally manipulated, the functional contributions of the observed gene-expression changes remain unresolved. Targeted loss- and gain-of-function experiments could establish whether changes in *Sp4*, *Cacna1g* and *C4b* contribute to subsequent circuit and behavioral alterations, and whether the P21 *Sp4* increase is compensatory or detrimental. We also did not test whether SCZ-associated variants modified the response to ELS; combining genetic perturbations with ELS would allow direct tests of gene– environment interactions. Second, we measured transcript abundance; whether these differences were accompanied by changes in protein abundance, activity or circuit function remains to be determined. Third, LBN was applied at the litter level, limiting independent replication to the number of litters represented. With four animals per condition at each age, several effects detected in stratified analyses did not reach significance in per-animal comparisons. Fourth, each animal was sampled at a single age, so the developmental trajectories reflected cross-sectional comparisons rather than changes tracked within individual animals. Finally, with two animals per sex per condition and age, the study was not powered to resolve sex-dependent responses.

Overall, our findings showed that ELS altered the developmental expression of genes implicated in SCZ, raising the possibility that environmental adversity contributes to vulnerability in part by perturbing genes that also confer substantial genetic risk.

## Methods

### Mice

All procedures followed National Institutes of Health guidelines and were approved by the Boston University Institutional Animal Care and Use Committee (protocol 2018-539). C57BL/6J mice (The Jackson Laboratory, 000664; RRID:IMSR_JAX:000664) were used for all experiments. Only first-time (primiparous) dams were used, and timed breeding pairs were established so that all dams received identical male exposure and equivalent single-housing durations before parturition. Mice were group-housed under a 12-h light–dark cycle (lights on at 07:00) with food and water available ad libitum, and offspring remained with the dam until weaning at postnatal day (P) 21. Both male and female offspring were studied.

### Limited bedding and nesting

Early-life stress (ELS) was induced with the limited bedding and nesting paradigm^12^. Pregnant dams were singly housed before parturition. On P2, the dam and litter were transferred to a cage fitted with a stainless-steel perforated platform positioned approximately 2.5 cm above the floor to separate the animals from the bedding and were provided one half of a Nestlet square (Ancare, NES3600) as nesting material. Cages were left undisturbed until P6 for a brief cage cleaning and then again undisturbed until P9, when the platform was removed and standard housing resumed; maternal behavior was observed twice daily from P2 to P9. Standard-reared (SR) control litters were housed conventionally. Pups were weaned at P21 and group-housed thereafter.

For molecular profiling, whole brains were collected at P10, P21, P40–45 and P90–95 (referred to throughout as P10, P21, P40 and P90) from a balanced cohort of 32 mice: 4 SR and 4 ELS mice at each age, comprising 2 males and 2 females per age and condition (16 SR and 16 ELS; 16 male and 16 female overall), obtained from 10 litters (5 SR, 5 ELS).

### Tissue preparation

Brains for multiplexed fluorescence in situ hybridization (mFISH) were extracted fresh, embedded in O.C.T. compound (Tissue-Tek, 4583), flash-frozen on dry ice and stored at −80 °C. Sagittal sections were cut at 16 µm on a cryostat (Leica CM1800) at −16 to −19 °C, mounted onto Superfrost Plus slides (Fisher, 1255015) and processed within one month of collection.

### Multiplexed fluorescence in situ hybridization

mFISH was performed with the RNAscope HiPlex12 Reagent Kit (488/550/650/750) v2 (Advanced Cell Diagnostics, 324409) according to the manufacturer’s instructions. The 24-plex panel comprised ten SCHEMA schizophrenia-risk genes (*Cacna1g*, *Grin2a*, *Gria3*, *Trio*, *Sp4*, *Rb1cc1*, *Setd1a*, *Xpo7*, *Cul1* and *Herc1*), the complement gene *C4b*, three activity and synaptic-scaffold genes (*Fos*, *Npas4* and *Gphn*) and ten cell-type markers (*Slc17a7* (*Vglut1*), *Slc17a6* (*Vglut2*), *Slc32a1*, *Pvalb*, *Th*, *Chat*, *S100b*, *Olig1*, *Olig2* and *Cx3cr1*); probe catalog numbers are listed in Supplementary Table 1. Transcripts were detected over six rounds of hybridization, imaging and fluorophore cleavage. Nuclei were counterstained with RNAscope DAPI (320858), sections were mounted in ProLong Diamond Antifade (Thermo Fisher, P36961), and coverslips were removed between rounds with saline–sodium citrate buffer.

### Imaging

Images were acquired on a Nikon Ti2-E microscope with a Yokogawa spinning-disk head and a 40×/1.15-NA water-immersion objective (Nikon CFI Apo LWD), using 405-, 488-, 561-, 640- and 750-nm laser lines. Whole sections were tiled with autofocus applied every other tile, and identical acquisition settings were used across specimens.

### Puncta detection and atlas registration

FISH puncta were detected and counted per cell using Big-FISH^16^, producing a per-cell transcript-count matrix that served as the input for all downstream analysis. DAPI images were registered to the Allen Mouse Brain Atlas with the QUINT workflow^17^: sections were assigned to a sagittal atlas plane in QuickNII^37^ and non-linearly warped in VisuAlign^37^, cells were segmented from the DAPI channel in Ilastik^38^, and Nutil combined the alignment and segmentation to assign each cell atlas coordinates and a brain-region label^39^.

### Cell-type classification

Cells were classified by a single-pass dominant-marker rule applied to the ten cell-type markers. A cell was assigned to the cell type of its most-expressed marker when that marker had at least one transcript and was at least twice as abundant as the next-highest marker (with the runner-up floored at 0.5 transcripts for the ratio), or when the runner-up was zero. Cells that failed this test but whose two most-abundant markers belonged to the same lineage were sub-classified by the lineage-defining marker: within the GABAergic lineage (*Pvalb*, *Slc32a1*), *Pvalb* ≥ 1 transcript defined PV interneurons and the remainder inhibitory neurons; within the excitatory lineage (*Vglut1*, *Vglut2*), cells were split by the more abundant marker; and within the oligodendrocyte lineage (*Olig1*, *Olig2*), *Olig1* ≥ *Olig2* defined oligodendrocytes and the remainder oligodendrocyte precursor cells. Cells whose two dominant markers spanned different lineages were left unclassified. This procedure classified 1,288,350 of 1,907,152 segmented cells (67.6%) into ten types: *Vglut1*⁺ excitatory, *Vglut2*⁺ excitatory, inhibitory, PV interneuron, Th⁺, cholinergic, astrocyte/glia, microglia, oligodendrocyte and oligodendrocyte precursor cells (OPCs). Cells were assigned to ten regions (cerebellum, cortex, fiber tracts, hippocampus, hypothalamus, mid/hindbrain and medulla, olfactory areas, striatum and pallidum, thalamus and ventricular system). Ventricular-system strata are not shown because isolated cells detached during mounting could not be distinguished from cells lining the ventricles.

### Differential expression

All differential-expression analyses treated the mouse as the biological replicate (n = 4 per condition per age). For each animal, puncta were summed by region × cell type (pseudobulk), and strata with fewer than ten cells were excluded. The library size of each stratum, its total puncta across the 24 panel genes, entered every model as an offset. Counts were modeled with negative-binomial (NB2) generalized linear mixed models in glmmTMB^40^. For the whole-brain contrasts, each gene was modeled as count ∼ condition × age + region + cell type + sex + offset(log library size), with random intercepts for animal and litter (litter alone if the combined model did not converge). ELS − SR contrasts at each age were obtained from estimated marginal means (emmeans), converted to log₂FC and Benjamini–Hochberg (BH)-corrected across the 11 × 4 = 44 gene × age family. Whether the ELS effect depended on age was tested per gene by a likelihood-ratio test comparing this model with the model without the condition × age term, both with a random intercept for animal, BH-corrected across the 11 genes. Region- and cell-type-resolved effects with ages pooled were fit within each region (cell type, age and sex as covariates) or within each cell type (region, age and sex as covariates) and BH-corrected per gene across strata. Effects for each region × cell type stratum at each age were fit per stratum with condition as the only fixed effect and BH-corrected per gene and age across strata. Region-level P21 effects and hypothalamus-only effects at each age were fit on cells that passed an off-lineage filter, which excluded any classified cell with two or more puncta of a cell-type marker belonging to a different lineage; hypothalamic models included cell type and sex as covariates, and P values were BH-corrected across the 11 risk-panel genes × 4 ages among fits that converged. Activity and scaffold markers (*Fos*, *Npas4*, *Gphn*) were modeled separately at each age (count ∼ condition + region + cell type + sex + offset(log library size), random intercepts for animal and litter, litter alone if the combined model did not converge) on units excluding unclassified cells and the ventricular system; P values for the condition term were BH-corrected within each marker across the four ages. Their stratum-level effects were BH-corrected within each marker across all age × region × cell type strata.

### Spillover correction, per-animal quantification and cross-age associations

To control for optical spillover of puncta between adjacent cells, we computed for each hypothalamic cell at P21 the mean count of the gene among its 15 nearest neighbors (all cell types; within each animal × subregion), z-scored this neighborhood term within each cell type and added it as a covariate to cell-level NB GLMMs (count ∼ condition + neighborhood + offset(log library size), random intercepts for animal and litter), requiring at least 40 cells and three animals per cell type; P values were BH-corrected across the gene × cell type family. Per-animal quantification used puncta of the target gene per 1,000 panel puncta in each animal (ratio of sums over cells, for animals with at least ten classified cells in the stratum), modeled with a linear mixed model with litter as a random intercept (lme4/lmerTest); P values are two-sided Satterthwaite tests of the condition term. Representative images show raw fluorescence with uniform brightness and contrast adjustments applied identically to SR and ELS. For cross-age associations, the ELS effect (log₂FC) of each of 14 panel genes was taken for every region × cell type stratum present at both P10 and P21, and Spearman correlations were computed between every P10 gene and every P21 gene across strata (pairs with fewer than six strata were not tested). All correlations were BH-corrected once as a single family. Throughout, ‘predictor’ refers to the earlier (P10) measurement. A UMAP embedding of classified cells was computed with uwot (n_neighbors = 15, min_dist = 0.1) from min–max-scaled puncta counts of 21 panel genes (ten cell-type markers, ten SCHEMA genes and *C4b*) for the 1,163,386 classified cells with at least 6 of the 24 panel genes detected; 30,000 randomly selected cells are shown.

### Statistics and reproducibility

No statistical method was used to predetermine sample size, but our sample sizes are comparable to those of previous single-cell spatial-transcriptomic studies. Rearing condition was assigned at the litter level because LBN is a manipulation of the dam and litter, and both sexes were included within each age and condition. The molecular pipeline was automated and applied identically to all samples. Cells without an assigned brain region were excluded from all models. No animals were excluded. All statistical tests were two-sided. Models with sex as a covariate estimated a single ELS effect across sexes; with two animals per sex per condition and age, the study was not powered to detect sex differences. Data are presented as mean ± s.e.m. unless otherwise noted, and n refers to the number of mice. Significance is denoted as P < 0.05 (*), P < 0.01 (**), P < 0.001 (***), and P ≥ 0.05 (ns). Per-animal comparisons use unadjusted P and show ns; other panels use Benjamini–Hochberg-adjusted P, as stated in each legend, and leave non-significant positions blank. Analyses were performed in R (v.4.5.1).

## Data availability

The data supporting the findings of this study can be explored through an interactive application upon publication.

## Code availability

Analysis code will be made available upon publication.

## Acknowledgments

We thank M. Sanchez-Carbonell for reading the manuscript and sharing her thoughts, and J. Chen and T. Blute for microscope access and assistance.

## Author contributions

A.B. and A.C.-M. conceived the study and designed the experiments. A.B. performed the experiments, developed the analysis pipeline, performed multiplexed fluorescence in situ hybridization (mFISH) staining, analyzed the data and prepared the figures. A.B. and A.C.-M. wrote the manuscript. R.A.P. contributed to the conceptual development of the study. R.A.P. and L.F. assisted with troubleshooting the mFISH experiments. M.B. performed preliminary mFISH staining. M.S. assisted with the early-life stress (limited bedding and nesting) paradigm. A.C.-M. supervised the project and acquired funding. All authors reviewed the manuscript.

## Supplementary figures

**Supplementary Fig. 1.**
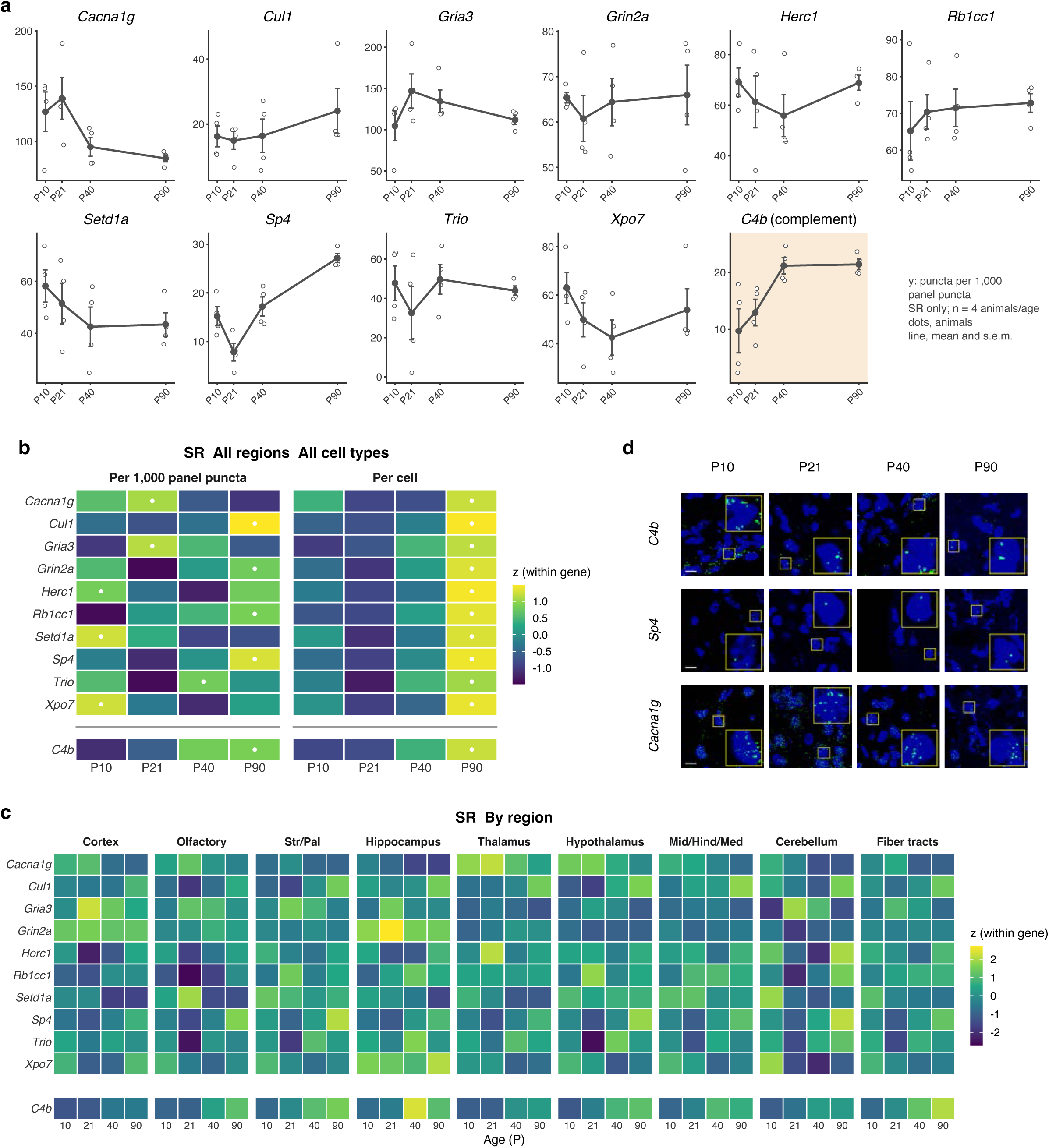
Developmental expression of risk-panel genes under standard rearing. a, Expression of the ten SCHEMA genes and *C4b* (shaded) in SR animals from P10 to P90, pooled across regions and cell types. Each gene is plotted on its own y axis. b, The SR means in a, z-scored within each gene, normalized to total panel puncta (left) or per cell (right). Dots mark the age of peak expression. Total panel puncta per cell was highest at P90, so per-cell values for all genes peak at that age. c, As in b (left), resolved by region; z-scores are computed within each gene across all regions and ages. d, Representative SR images at each age for *C4b* (thalamus), *Sp4* (thalamus) and *Cacna1g* (hypothalamus): gene (green) and DAPI (blue), with one display range per gene across ages; callouts, 3× magnification of a typical positive cell (yellow box; at least four puncta for *C4b* and *Cacna1g*, at least one for *Sp4*). Scale bar, 10.1 µm (P10 images; all images share one scale). Expression is puncta per 1,000 panel puncta. Points, individual animals; lines and error bars, mean ± s.e.m. n = 4 SR animals per age (16 animals, 5 litters).

**Supplementary Fig. 2.**
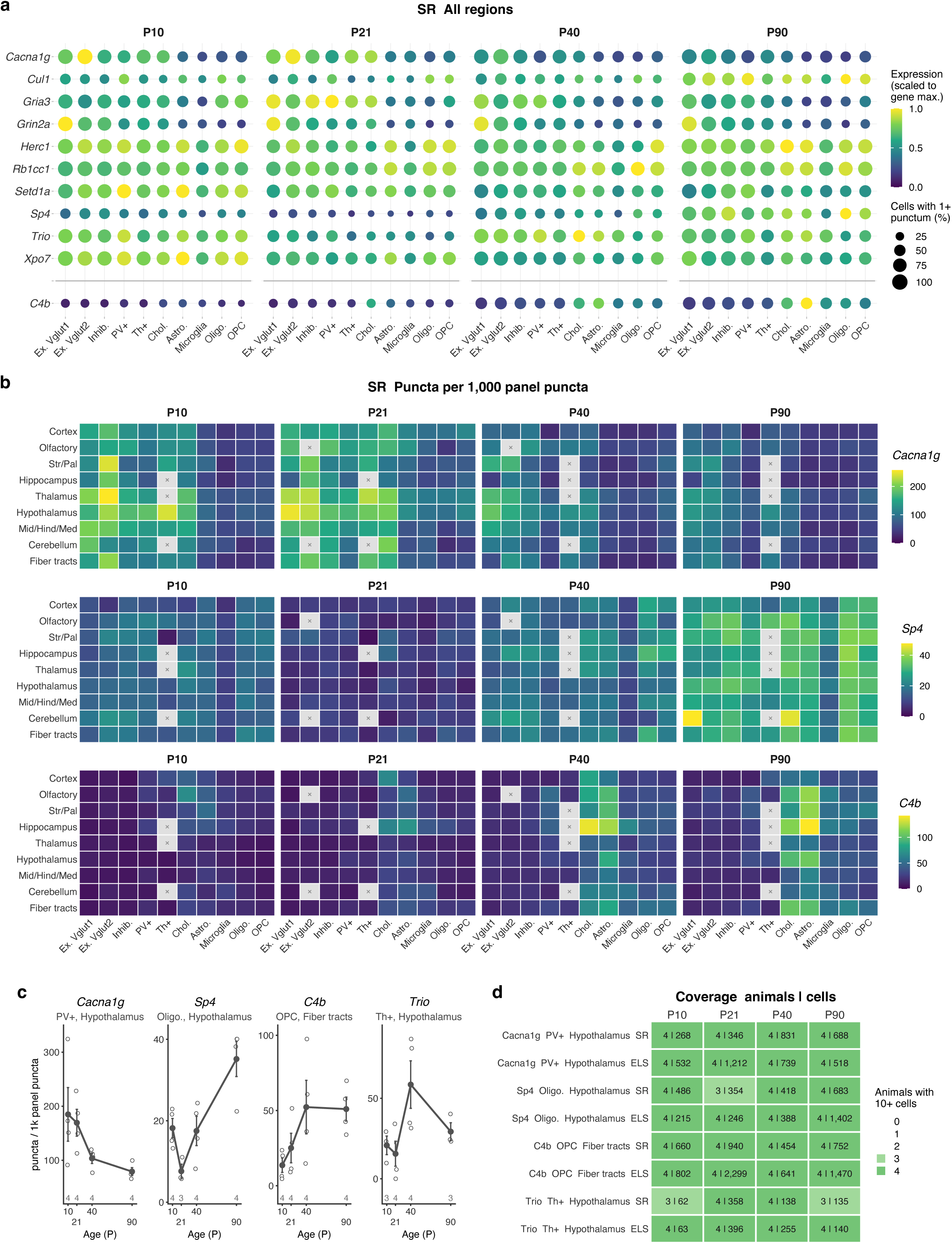
Cell-type and regional expression of risk-panel genes under standard rearing. a, Expression of the ten SCHEMA genes and *C4b* across cell types at each age, pooled across regions. Color, expression scaled to the maximum for each gene; dot size, percentage of cells with at least one punctum. b, Expression of *Cacna1g*, *Sp4* and *C4b* by region and cell type at each age, with one color scale per gene. Crosses mark strata sampled in fewer than two animals. c, Developmental trajectories of *Cacna1g* in hypothalamic PV⁺ interneurons, *Sp4* in hypothalamic oligodendrocytes, *C4b* in fiber-tract OPCs and *Trio* in hypothalamic Th⁺ cells. Points, individual animals; lines and error bars, mean ± s.e.m.; numbers, animals per age. d, Animals contributing at least ten cells, and total cells, for each population in c by condition and age. Expression is puncta per 1,000 panel puncta, computed per animal from strata with at least ten cells. a–c, SR animals only (n = 4 per age; 16 animals, 5 litters) unless noted; in b some strata have 2 or 3 animals, and numbers in c give animals per age.

## Supplementary table

**Supplementary Table 1:**
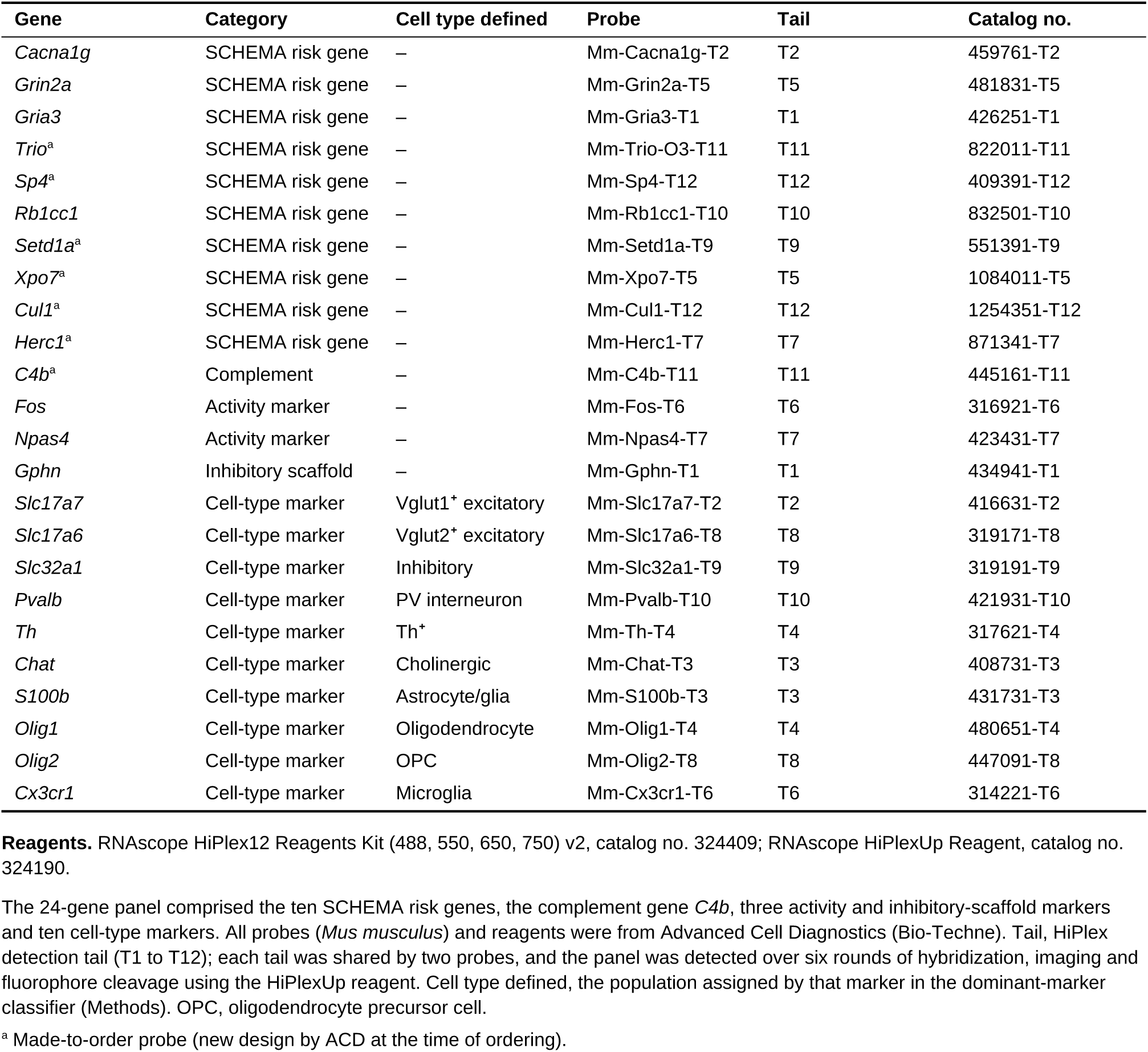
RNAscope HiPlex probe catalog (mouse). The 24 panel genes (10 SCHEMA risk genes, complement *C4b*, three activity/synaptic markers, ten cell-type markers) with ACD probe names and catalog numbers.

